# Separation of the effects of small intestinal microbiome-diet interactions on human gut hormone secretion

**DOI:** 10.1101/2022.09.25.509333

**Authors:** Sara C. Di Rienzi, Heather A. Danhof, Juan Huerta, Robert A. Britton

## Abstract

Microbial regulation of gut hormones is a potential mechanism by which the gut microbiome acts on systemic physiology. However, there are limited systems that permit study of how small intestinal microbes and diet modulate gut hormone secretion. Here we present the platform **C**ulturing and **A**pplication of **M**icrobes on **I**ntestinal **O**rganoids (CAMIO) and demonstrate its usage in studying the effects of diet and microbes on gut hormones. We validate that CAMIO supports long-term cultivation of a small intestinal microbiome in different dietary sugars and show that CAMIO permits measurement of gut hormones released from jejunal organoids in response to products of the small intestinal communities. In doing so, we observe differential secretion of ghrelin, PP, and PYY according to whether the microbial communities were grown in glucose-fructose versus sucrose or trehalose. We expect CAMIO to be useful in mechanistically understanding how diet and microbes collectively regulate gut hormones.

## Introduction

In his efforts to teach medicine, Hippocrates is attributed to the statement: “All disease starts in the gut” and he further advocated that proper nutrition is the foundation of health. Hippocrates made these statements when biology and medicine were still in their infancy and well before there was knowledge of how information from the gut traveled to other parts of the body. Today we understand that one of the main ways the presence of food in the gut is communicated to the brain and the rest of the body is through molecules and peptides called hormones (Bellono et al., 2017; Kaelberer et al., 2018; Mace et al., 2015; McCauley et al., 2020).

Many of these gut hormones are produced in enteroendocrine cells (EECs) of the small intestine where they rapidly sense consumed food and differentially respond (Beumer et al., 2020; Fazio Coles et al., 2020; Gribble and Reimann, 2016, 2017). For example, glucose consumption suppresses secretion of the hormone ghrelin (Djurhuus et al., 2002; Parker et al., 2005; Sinagoga et al., 2018; Tschöp et al., 2000) and enhances peptide tyrosine tyrosine (PYY) (Adrian et al., 1985; Gerspach et al., 2011; Steinert et al., 2011) and pancreatic polypeptide (PP) secretion (Aragón et al., 2015; Sive et al., 1979). Fructose consumption on the other hand, causes less potent suppression of ghrelin secretion (Teff et al., 2004; Van Name et al., 2015), and lower or similar release of PYY than glucose (Kuhre et al., 2014; Lindqvist et al., 2008). These hormones are communicated to the brain and subsequently modulate feeding behavior (Asakawa et al., 2003; Banks et al., 2002; Date et al., 2002; Kaelberer et al., 2018; Nonaka et al., 2003). PYY and PP have a satiety effect whereas ghrelin produces the hunger sensation (Batterham et al., 2002; Nakazato et al., 2001; Ueno et al., 1999).

As complex as this system of gut hormonal signals is, there is a further level of complexity that regulates the response to food, the gut microbiome. In recent years, it has been discovered that gut microbes regulate the secretion of a variety of gut hormones, including PYY and ghrelin (Engevik et al., 2021; Martin et al., 2019; Perry et al., 2016; Tomaro-Duchesneau et al., 2020; Yano et al., 2015). However, it is also well recognized that gut microbes are affected by and affect diet (David et al., 2014; Di Rienzi and Britton, 2020; Kolodziejczyk et al., 2019; Makki et al., 2018), which regulates gut hormones. This sets up a complex tri-cornered paradigm of gut hormonal secretion.

Approaches are thereby needed to understand the contribution of diet and the small intestinal gut microbiome to gut hormone secretion. Given the expense and size needed for a properly controlled clinical study and the limitations of animal models in their relevance to human health (Nguyen et al., 2015), *in vitro* models are an attractive alternative. These models consist of bioreactors modeling diet-microbe interactions and cell culture models modeling host-microbe interactions.

For bioreactors, there are large-scale dynamic bioreactors whose primary goal is to model digestion (Havenaar et al., 2013; Minekus et al., 1995, 1999; Molly et al., 1993; Stolaki et al., 2019). These consist of multiple vessels that model the different regions of the intestine. Media and secretions coming into the different vessels are dynamically controlled so to regulate the pH or release food at time intervals. Work by Stolaki and colleagues (Stolaki et al., 2019) was successful in using this type of system to cultivate human ileostomy samples for 14 days. These models are limited in that they are very large and thus performing replicates is difficult. In response to this limitation, small volume bioreactors have been designed which simplify the reactor system and reduce the working volume, thereby facilitating a higher number of replicates. The work by Cieplak et al (Cieplak et al., 2018) is particularly notable in that they added to the ileum compartment 7 small intestinal strains. While culturing was short term, this work marked the first attempt at culturing small intestinal communities in bioreactors.

For cell culture, there are models that try to recreate the architecture of the intestine by utilizing a scaffold onto which cells adhere (Chen et al., 2017; Costello et al., 2014; Wang et al., 2017; Zhou et al., 2021). Other models utilize a simpler transwell or monolayer format but focus on host-microbe interactions (Calatayud et al., 2019; Roodsant et al., 2020). Some of these models utilize mixtures of different cell types. For example, Calatayud and colleagues (Calatayud et al., 2019) used a triculture of Caco-2, HT29-MTX, and THP-1 cells on transwells and tested the response of these cells to a community of 8 small intestinal strains. Alternatively, human intestinal organoid cultures can be used, which recapitulate all the cell types of the tissue from which they were generated from thereby increasing translation to human health (Date and Sato, 2015; Sato et al., 2009). Overall, these existing methods have made progress in modeling the small intestine but are lacking in their ability to measure secreted gut hormones in a biologically relevant context.

Here we report a platform, which improves upon these existing approaches. The platform supports long-term growth of small intestinal microbes in continuous culture bioreactors, easy replication, study of diet-microbe interactions, and incorporates a recent intestinal organoid model of gut hormone secretion (Chang-Graham et al., 2019). We term this platform **C**ulturing and **A**pplication of **M**icrobes on **I**ntestinal **O**rganoids (CAMIO).

## Results

### CAMIO overview

CAMIO has two main components: A) 15 ml volume bioreactors utilizing a fully defined medium designed to simulate the small intestinal environment, including having all 4 primary human bile acids and a nutrient concentration and flow rate like that of the small intestine; B) Human small intestinal (jejunal) organoids that have been genetically engineered to be enriched in enteroendocrine cells (Chang-Graham et al., 2019) (see Methods). These organoids allow for measurement of gut hormones in response to stimuli in a physiological relevant context.

To use CAMIO, dietary components of interest are selected and incorporated into the bioreactor medium (**Figure 1**). Next, microbes or a microbial community are inoculated into bioreactors containing the medium and cultured with continuous flow in the bioreactors for a length of time or number of generations of interest (**Figure 1**). During this time, the community is sampled to monitor how microbial density and composition are affected by the dietary component. Lastly, experimental supernatants from the bioreactor microbial communities are prepared (cell-free, pH neutralized) and applied to the enteroendocrine enriched human intestinal organoids for measurement of gut hormone release (**Figure 1**).

**Figure 1:**
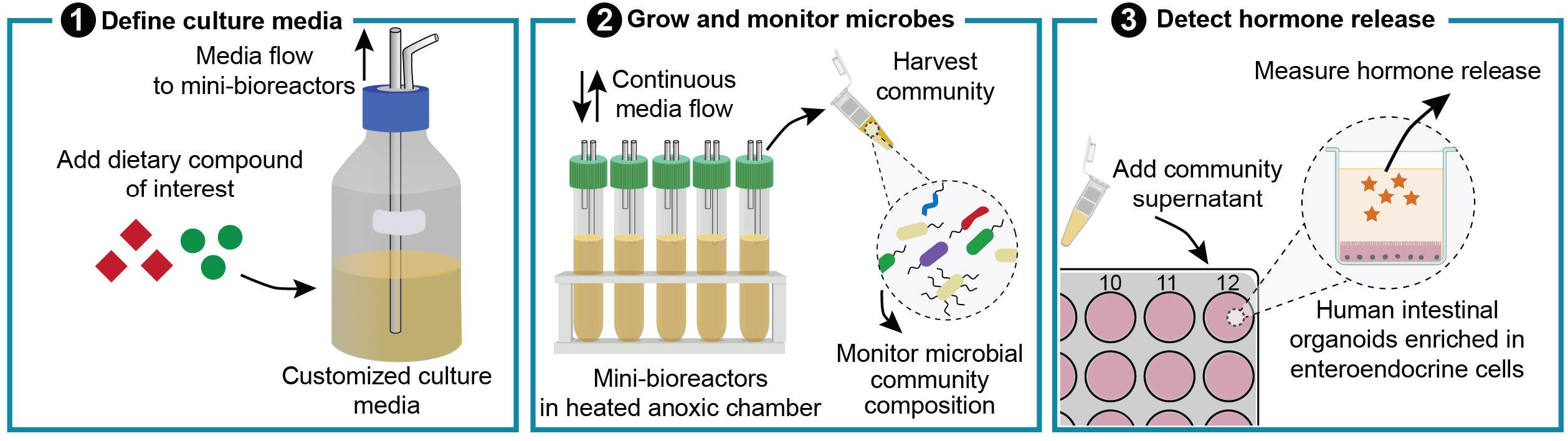
Overview of CAMIO. To use CAMIO, first, dietary compounds of interest are selected and incorporated into a medium that simulates the small intestinal environment. Second, small intestinal microbes are inoculated into mini-bioreactors, and flow is sustained in the bioreactors for a length of time of interest. The microbial community composition is monitored by sequencing samples taken from the bioreactors. Third, cell-free, neutralized supernatants are prepared from the bioreactor microbial communities and applied to human intestinal organoids enriched in enteroendocrine cells.

To validate the functionality of CAMIO, we had three main goals: demonstrate that CAMIO can 1) grow a small intestinal community, 2) study the effects of diet on microbes, and 3) measure the secretion of gut hormones in response to diet and microbes.

### CAMIO enables long-term growth of a simple small intestinal community

To test if CAMIO permits growth of small intestinal microbes, we selected human microbes (**Table 1**) that are largely predicted to be prevalent, abundant, or of interest to the function of the human small intestine (Leite et al., 2019; Seekatz et al., 2019; Sundin et al., 2017; Villmones et al., 2022; Wang et al., 2005). We therefore utilized a six-member community comprised of human-derived *Prevotella histicola, Lacticaseibacillus casei, Limosilactobacillus reuteri, Streptococcus mitis, Streptococcus salivarius*, and *Veillonella atypica* (**Table 1**). The medium was designed such that our dietary component of interest was sucrose, and it represented 87% of the macromolecules – carbohydrates, protein, and fat.

**Table 1.**
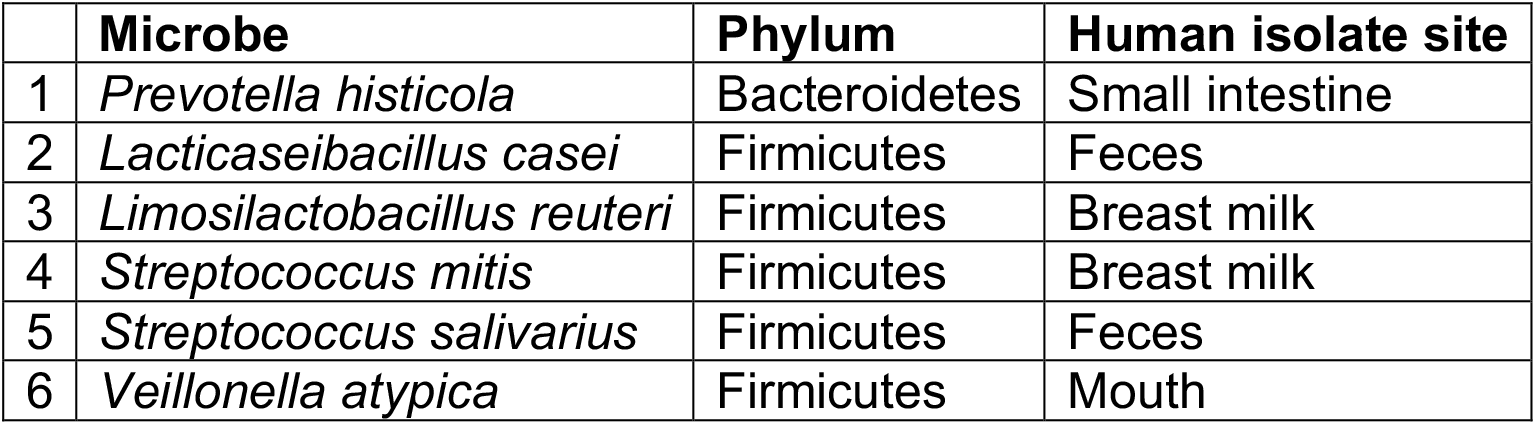
Microbial strains used to model the small intestinal microbiota.

The small intestinal microbial strains and sucrose bioreactor media were inoculated into twelve reactors in sets of four, with each set supplied a different media bottle (see Methods). Microbial community composition was followed by sampling the flow-through of the reactors. The microbial communities of the three different sets were compared by assessing their beta-diversity as measured by a Bray-Curtis distance metric. Utilizing principal coordinate analysis and PERMANOVA, we note that while differential communities can be observed at generation 0 before flow is initiated on the reactors, by generation 50 the communities have significantly converged (**Supp Fig 1A**).

More detailed analysis revealed that the communities converge on a predominantly three-membered community of *L. casei, V. atypica*, and *S. salivarius* and that *P. histicola* was lost (**Figure 2**). The communities held a density of 10^8^ cells/ml during this time (**Suppl Fig 1B**). From these observations, we conclude that CAMIO does permit culturing over multiple generations of a simple small intestinal community.

**Figure 2:**
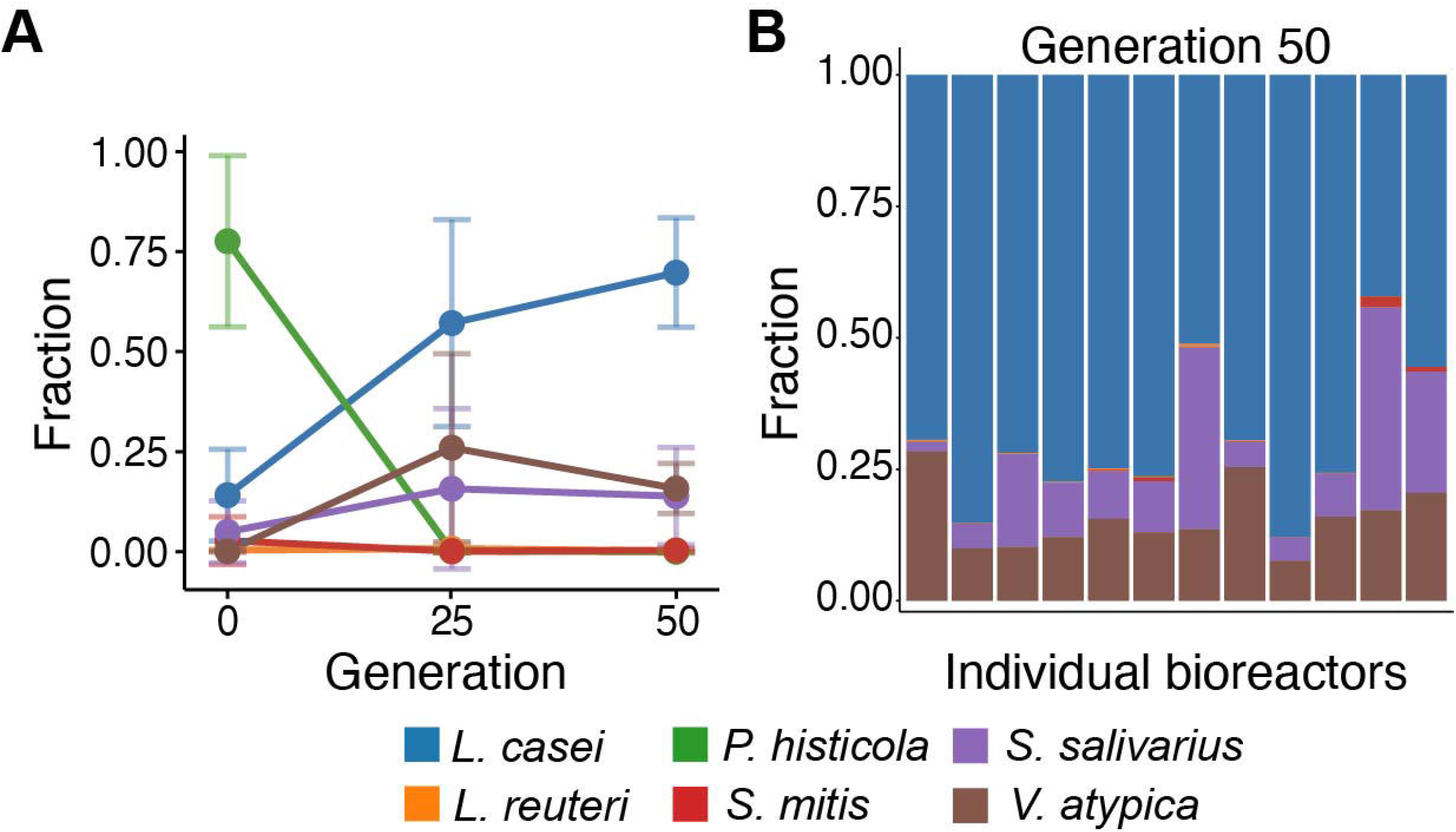
Small intestinal bioreactor communities growing on sucrose media for 50 generations. **A)** Mean relative abundance of each microbe across all bioreactors from generations 0 to 50. **B)** Relative abundance of each microbe in each bioreactor at generation 50. Generations are averaged across reactors. See Supplemental Figures 1 and 2 for further details of the communities.

### CAMIO enables study of the effect of diet on microbes

To test if CAMIO could be used to study how diet affects small intestinal microbes, we continued culturing the same reactors but replaced the sucrose in the media to either a mixture of 45% glucose, 55% fructose (here referred to as high fructose corn syrup, HFCS) or trehalose or kept it as sucrose and continued culturing the communities for another 100 generations. Over this time, the communities remained at a density of 10^8^ cells/ml (**Suppl Fig 1B**). In the sucrose reactors, the communities appeared to have a diversity similar to that at generation 50 (**Figure 3A, Suppl Fig 2**). The HFCS and trehalose reactors, however, appeared to have an increased abundance of *V. atypica* and a reduction of *S. salivarius* (**Figure 3A**). We also observed cycling between *V. atypica* and *L. casei* (**Suppl Fig 2**). Statistical analysis of the communities before and after the change in sugars, confirmed that *S. salivarius* was decreased significantly in the trehalose reactors (**Figure 3B**) while *V. atypica* was increased (**Figure 3C**). Changes in the HFCS communities did not reach statistical significance (**Figure 3B,C**).

**Figure 3:**
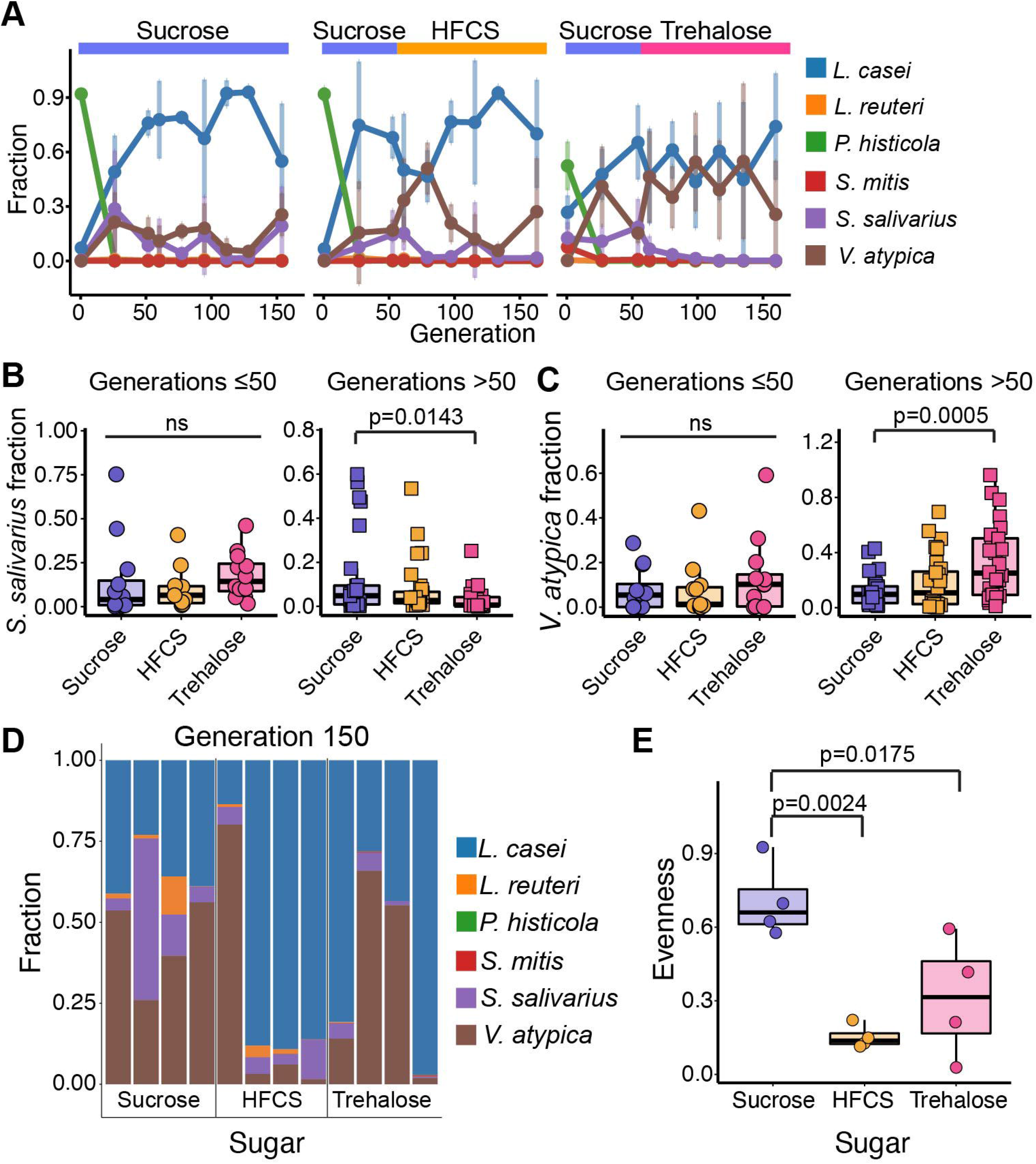
Small intestinal bioreactor communities after switching to HFCS or trehalose media for 100 additional generations. **A)** Mean relative abundance of each microbe across bioreactors by their sugar media from generations 0 to 150. Generations are averaged across reactors. Relative abundance of **B)** *S. salvarius* and **C)** *V. atypica* in reactors before (days 0, 3, and 6; generations ≤ 50) and after changing sugar media (days 7 to 18; generations > 50) as assessed from flow-through. **D)** Final community composition within the reactors as assessed by direct sampling of communities from the reactors. **E)** Species evenness of the reactors at the termination of the experiment as in D. See Supplemental Figures 1 and 2 for further details of the communities.

After 150 generations, we assessed the final microbial communities by sampling the reactors directly. The HFCS and trehalose reactors were either mostly *V. atypica* or *L. casei*, whereas the sucrose reactors had a more balanced composition among *V. atypica* and *L. casei* and the other strains (**Figure 3D**). Moreover, the microbial community structure was statistically less even in the HFCS and trehalose reactors (**Figure 3E**).

Additionally, CAMIO facilitates more in depth analyses of diet-microbe interactions. First, as CAMIO allows for time-course sampling of the bioreactors, how and when a dietary shift affects the microbial communities can be measured. While all communities had similar beta-diversity at generation 50 before the sugars were switched in the media (**Figure 4A, Supp Fig 1A**), we observe that at generation 75 the three sets of reactors were able to be differentiated by the sugar (**Figure 4A, Supp Fig 1A**). The communities’ beta-diversity became indistinguishable again until generation 125 at which we observe that the trehalose reactors separate from the sucrose and HFCS reactors (**Supp Fig 1A**).

**Figure 4:**
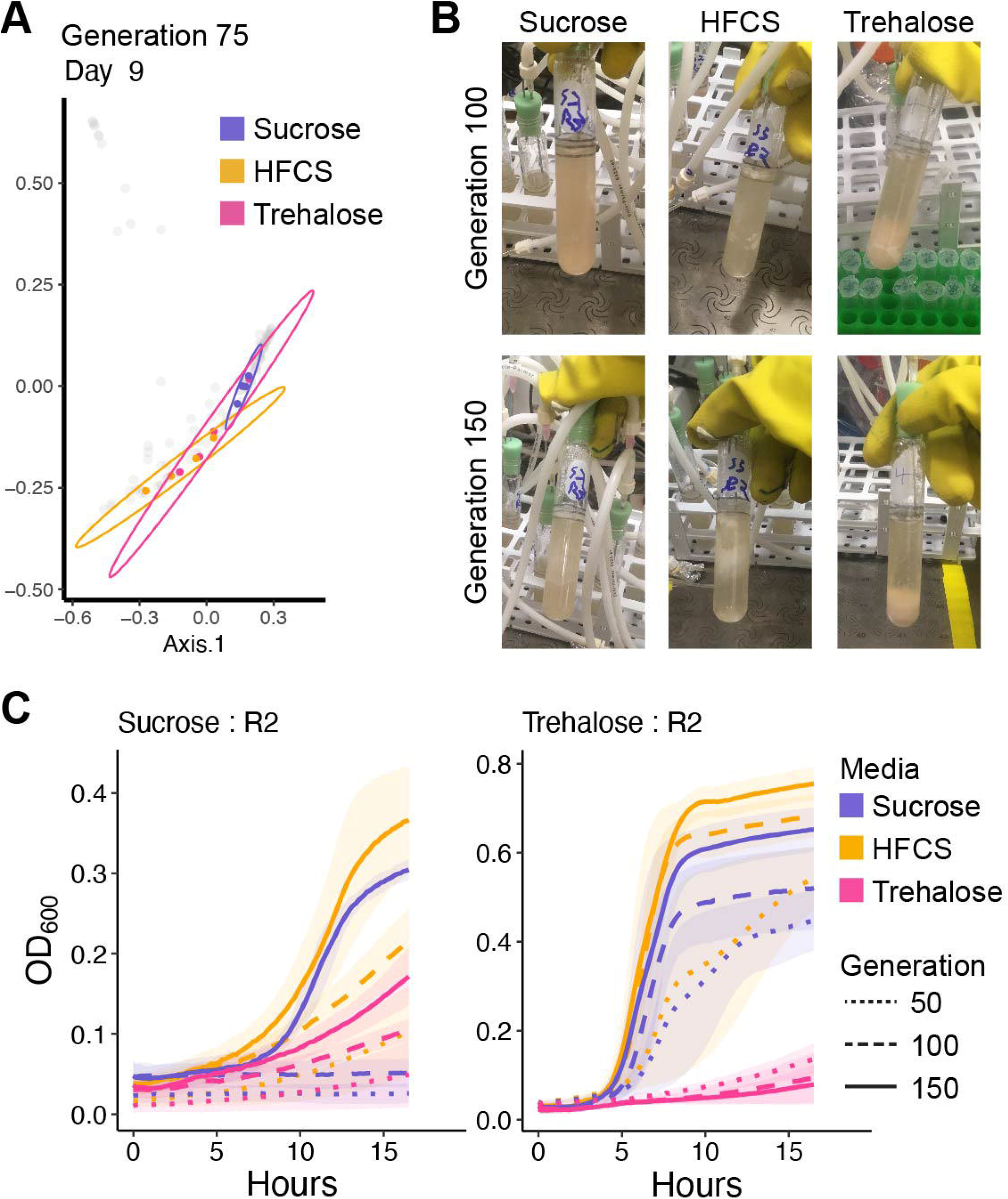
Small intestinal community composition, physical appearance, and growth adaptation resulting from growth in different sugars. **A)** PCoA as in Supplemental Figure 1 of the bioreactor communities at generation 75. **B)** Images of representative bioreactors in small intestinal media with each sugar at generations 100 and 150. **C)** Growth dynamics of generations 50, 100, and 150 from sucrose reactor 2 and trehalose reactor 2 when regrown in batch phase in each different sugar media. Growth dynamics of all reactors are shown in Supplemental Figure 3.

Second, CAMIO uses glass bioreactors, allowing for physical visualization of the microbial communities at all times. During our study, each sugar gave rise to a different culture appearance. The sucrose communities had a homogenous and slightly pink color, the trehalose communities were of a similar color but fell to the bottom of the tube when not stirred, and the HFCS communities were of a more yellow-green color and had a thick biofilm (**Figure 4B**). The biofilm adherent to the HFCS glass reactors was sequenced and identified to be *L. casei*.

Third, CAMIO can be used to isolate or study microbial adaptation to diet. As a demonstration of this use, we collected microbial communities from generations 50, 100, and 150 in each culture condition and then regrew the communities in batch culture in each of the three sugars. If adaptation had occurred, we expected the 150-generation community to grow better than the previous generations in one or more of the sugar media. Several of the bioreactor communities displayed such adaptation (**Suppl Fig 3**). For example, sucrose reactor 2 and trehalose reactor 2 generation 150 communities grew better on each of the different sugars than the previous generation communities on each of the different sugars (**Figure 4C, Suppl Fig 3**). The adaptation in trehalose reactor 2 differed from that in sucrose reactor 2; however, the trehalose reactor 2 generation 100 community grew worse than the trehalose reactor 2 generation 50 community (**Figure 4C, Suppl Fig 3**).

### CAMIO enables testing of diet and microbial products for human hormone secretion

Lastly, we asked if CAMIO allows for testing of how diet and microbe interactions alter gut hormone secretion. We tested media alone and cell-free, pH neutralized bioreactor supernatants on enteroendocrine cell-enriched organoid monolayers for 3 hours. We were able to measure C-peptide, ghrelin, GIP, GLP-1, MCP-1, PP, PYY, serotonin, and TNFα secreted from the organoids (**Suppl Fig 4**). Only ghrelin, PP, and PYY showed differences among the media and microbial supernatants. We observed that HFCS media alone led to greater secretion of PYY (**Figure 5A-C**). When normalized to media controls, microbial supernatants from the end of the bioreactor run from HFCS reactors led to greater secretion of ghrelin, and lower secretion of PP, and PYY (**Figure 5D-F**).

**Figure 5:**
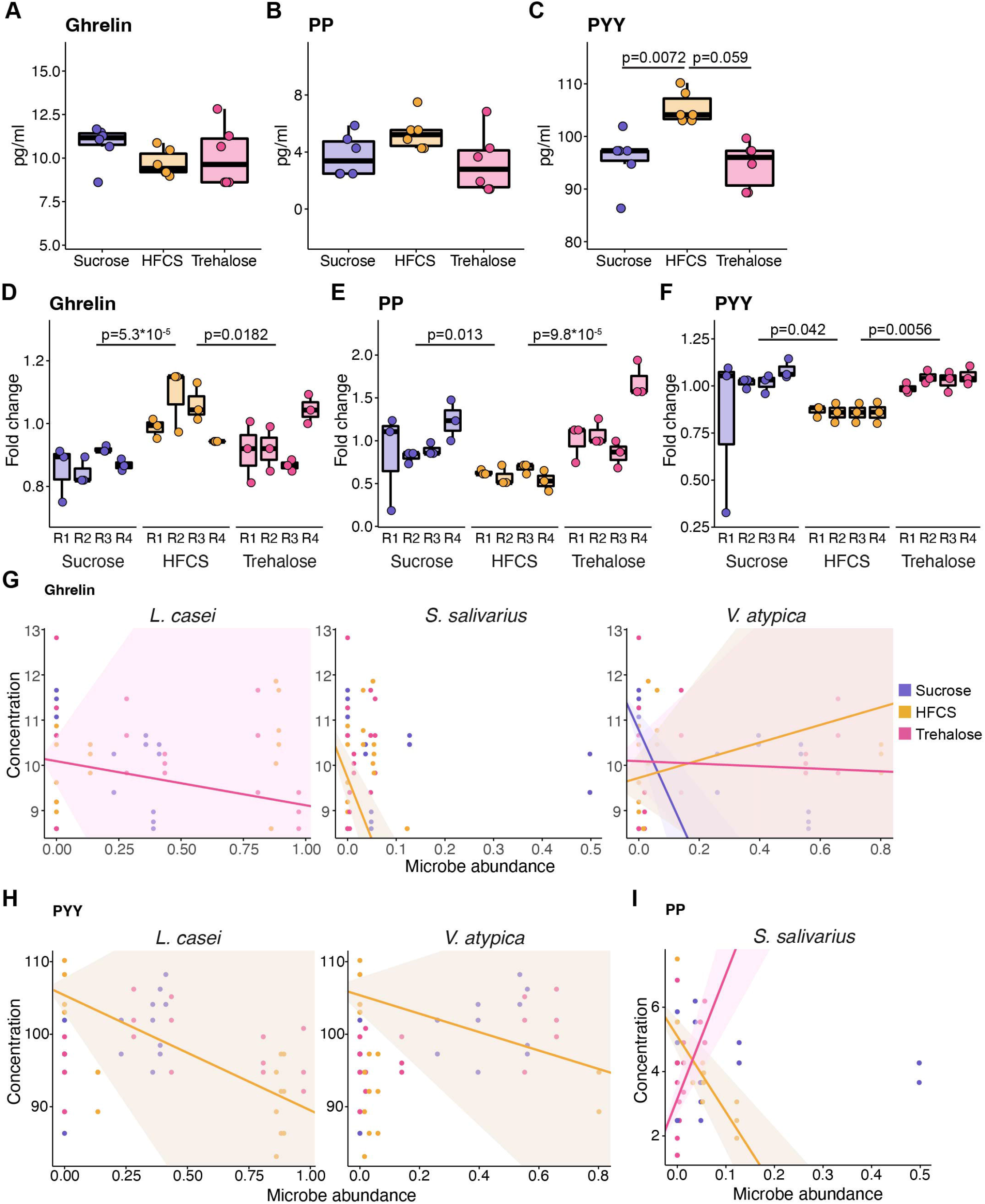
HFCS-grown microbial communities differentially regulate hunger and satiety hormones. Secreted **A)** ghrelin, **B)** PP, and **C)** PYY from enteroendocrine-enriched organoid monolayers treated with bioreactor media. Fold change over media control of **D)** ghrelin, **E)** PP, and **F)** PYY secreted from organoid monolayers treated with bioreactor communities on sucrose, HFCS, or trehalose media. Hormone concentration VS microbe abundance for **G)** ghrelin, **H)** PYY and **I)** PP. Lines are drawn when the microbe:sugar interaction was significant in a linear model. Shaded regions indicate the standard error of the fitted line. See Supplemental Figure 4 for all unnormalized hormone data.

Using the microbial abundance data from the bioreactors, we aimed to see if we could use linear modeling to predict which microbes were contributing to observed hormonal secretion. We started with a full model of all the microbes interacting with the media. Then we removed microbes one by one provided the removal didn’t affect the predictive ability of the model. In doing so, we were able to create the following simplified models: ghrelin as a function of *L. casei, S. salivarius*, and *V. atypica* interacting with the media; PP as a function of *S. salivarius* interacting with the media, and PYY as a function of *L. casei* and *V. atypica* interacting with the media.

Inspection of the models indicated which microbe interacting which media best explain the observed data. Ghrelin was overall best explained by *V. atypica*, which decreased ghrelin levels when grown on sucrose or, to a lesser extent, trehalose, and increased by *V. atypica* grown on HFCS (**Fig 5G, Suppl Table 1**). Decreased ghrelin levels were also associated with *S. salivarius* grown in HFCS and *L. casei* grown in trehalose (**Fig 5G, Suppl Table 1**). PYY levels could be predicted best as decreased by *L. casei* and *V. atypica* when grown in HFCS (**Fig 5H, Suppl Table 1**). Lastly, PP could be predicted as increased when *S. salivarius* was grown in trehalose and decreased when it was grown in HFCS (**Fig 5I, Suppl Table 1**).

## Discussion

The dimensionality of the gut ecosystem is infinite due to the complexity of the diet, gut microbes, and host cells. While existing methods can propagate in a biologically relevant context small intestinal microbes and allow for study of how specific dietary components affect microbes, they lack the ability to study how those microbes in turn alter gut hormone secretion. We aimed to fill this gap and make a system that can address each of these critical variables of human physiology. Our data presented here, demonstrates that CAMIO meets these goals.

### Observed results

Our specific investigation of six small intestinal strains led to microbial communities with *L. casei* and *V. atypica* as the most abundant strains. Only the sucrose reactors resulted in a more even community composition. When grown alone in the sucrose media, *L. casei* demonstrated a long lag phase that was not observed by the other streptococci and lactobacilli (**Suppl Fig 5A-D**). This lag may have suppressed its growth enough to permit increased growth of *S. salivarius. P. histicola* was lost in the bioreactors and grew poorly alone in all small intestinal media (**Suppl Fig 5E**). In contrast *V. atypica* was a dominant member of all bioreactor communities yet only grew somewhat well on the HFCS media alone (**Suppl Fig 5F**). This observation in conjunction with evidence of cycling between *L. casei* and *V. atypica* suggests these strains may have been cross feeding in the reactors. *V. atypica* is known to cross-feed with other microbes via lactate production (Scheiman et al., 2019). Therefore, it is likely that we have created a cross-feeding system between *L. casei* and *V. atypica*. In further support of this notion, we observed *V. atypica* increased in abundance in the trehalose reactors and *S. salivarius* decreased. *L. casei* was the only strain capable of growing well in trehalose media alone (**Suppl Fig 5**). Hence, we interpret these observations to imply the trehalose media is supporting the growth of *L. casei* which in turn supports the growth of *V. atypica*.

Our investigations on these six strains growing in sucrose, HFCS, or trehalose also led to the observation that HFCS microbial supernatants promoted higher secretion of ghrelin and lower secretion of PYY and PP. Secretion of ghrelin in response to HFCS/fructose VS sucrose/glucose has been studied in rats (Kisioglu and Nergiz-Unal, 2018) and humans (Teff et al., 2004). In both systems, HFCS led to a 1.05-to-1.5-fold increase in ghrelin, similar to our data. However, several other studies have reported no change in ghrelin plasma levels in response to sucrose VS HFCS in humans (Melanson et al., 2008).

Interestingly, sucrose and trehalose media alone reduced PYY secretion. One explanation for these differences is that HFCS is a mixture of the monosaccharides glucose and fructose while sucrose and trehalose are disaccharides. Therefore, perhaps for the media alone, the glucose in HFCS is rapidly or strongly stimulating PYY. Similarly, in the microbial supernatants, the glucose and perhaps also the fructose were likely consumed by the microbes and hence were not available to the organoids. These results further imply that the breaking of the disaccharide bond in sucrose and trehalose by the microbes or the organoid cells attenuates the organoid cell response to these sugars.

### Limitations and future improvements

CAMIO was designed as a first step in the study of the role of diet and microbes on gut hormone secretion. As such, there are several simplifications we used for this platform that could be improved upon. First, we used a simple static single-chambered bioreactor model. A dynamic bioreactor system that actively monitors and responds to pH and simulates the human feeding pattern would greatly enhance our ability to model the small intestinal environment. Additionally, the only host secretion we included in our media was bile. Other host secretions such as mucus and digestive enzymes could be included to more closely mimic host digestion. Second, here we assembled a small intestinal community from single strains. Utilizing a sample directly from the human small intestine would improve our study of the small intestinal microbiome and is also predicted to avoid the community composition from being dominated by one or two strains. It should be noted that the variation among replicate reactors that we observed is a known property of stochasticity in microbial community assembly in bioreactors (Auchtung et al., 2015; Oliphant et al., 2019). Third, the bioreactor style used here was one made of a tall glass tube. This configuration allowed for observational monitoring of the communities but made direct sampling from the bioreactors difficult without removal of the cork sealing the reactors. As such, we used flow-through sampling to monitor the community structure during the bioreactor run. If direct sampling is needed frequently, a bioreactor style like mini-bioreactor arrays (Auchtung et al., 2016) could be used. Lastly, we used a model of gut hormone secretion from an engineered jejunal organoid line (Chang-Graham et al., 2019). If hormone secretion from a non-engineered organoid line is needed, changes to the organoid growth media could be employed to enhance the abundance of enteroendocrine cells (Fujii et al., 2018; Villegas-Novoa et al., 2022; Zeve et al., 2022).

### Applications for CAMIO

We anticipate CAMIO being useful in several different approaches for studying diet-microbe-hormone or any diet-microbe-epithelial layer interaction. CAMIO would facilitate a forward screen for diet:microbe interactions that affect hormone production. These results could then be validated in an animal or clinical study. Conversely, CAMIO is equally suitable for follow up experiments to a clinical or animal study for the purpose of determining mechanism or further studying altered hormone secretion. Finally, CAMIO can also be used tangentially for the purpose of studying or creating diet-adapted microbial strains.

In summary, we have created a novel platform for exploring how gut hormones are regulated by diet and small intestinal microbes. We anticipate that further use and refinement of CAMIO will lead to insights into the role of the small intestinal microbiota in health and disease and ultimately assist in laying the foundation for novel treatments to cure systemic disease by targeting the gut ecosystem.

## Significance

Research has demonstrated that the gut microbiome is a potent effector of health and disease in body sites distal to the gut. Interactions with diet and regulation of gut hormones are postulated to mediate many of these effects. However, study of how diet and microbes affect physiology through gut hormones has been limited due to the lack of models to study the small intestine and small intestinal hormone secretion. Our platform CAMIO provides these tools allowing for study of how any dietary compound alters small intestinal microbiomes and their effect on hormone secretion. We demonstrate that CAMIO permits growth of small intestinal microbiomes in a defined diet of choice and study of how those dietary compounds alter microbial community composition. We specifically show that products from microbial communities grown in HFCS differentially promote secretion of ghrelin, PYY, and PP. The data generated by CAMIO thereby suggests that hunger and satiety can be modulated by interactions between the diet and microbiome and provides the methodology and platform for other investigations into how specific dietary compounds modulate gut hormone secretion through microbial products.

## Acknowledgements

We thank Jihwan Lee for creating Figure 1. We also thank Oleg Igoshin, Jenny Auchtung, Kylie Farrell, Susan Venable, and Torsten Seemann for their assistance, advice, and critical comments. SCD was supported by a fellowship from the NLM Training Program in Biomedical Informatics and Data Science (T15LM007093). HAD was supported by a fellowship from the National Institute of Allergy and Infectious Diseases (NIAID) grant F32AI136404. RAB was supported by NIH grants R01AI123278, R33AI121522, and U01AI124290.

## Author contributions

Conceptualization: SCD, RAB. Data curation: SCD, JH. Formal Analysis: SCD. Funding acquisition: SCD, HAD, RAB. Investigation: SCD, HAD, JH. Methodology: SCD, HAD. Project administration: RAB. Resources: RAB. Software: SCD. Supervision: RAB. Validation: SCD, HAD. Visualization: SCD. Writing – original draft: SCD. Writing – review & editing: SCD, HAD, RAB.

## Methods

### Bioreactors

The bioreactors used were a combination of designs developed by the Dunham (Miller et al., 2013) and Britton labs (Auchtung et al., 2016). In brief, media was contained in 2 liter bottles attached with tubing that ultimately ran through a peristaltic pump (series 205S, Watson-Marlow, USA) as described by Auchtung and colleagues (Auchtung et al., 2016). The media then flowed through an 18-gauge, 3-inch needle (N183, Air-Tite Vet Premium Hypodermic Needles, USA) set in a size 0 silicone cork (7752-15, Cole-Parmer, USA) in a 20 × 125 mm glass reactor tube (9826, Pyrex, USA). Media within the glass reactor tube was stirred by a small magnet and the reactors were set in a rack on a magnetic stir plate (MIXDrive60, 2mag-USA, USA). The media flowed out of the reactor by negative pressure via a 16-gauge, 3-inch needle (N163D, Air-Tite Vet Premium Hypodermic Needles, USA) positioned at the level within the glass reactor such that the media level was at 15 mls. Negative pressure within the tubes was obtained by a second set of peristaltic pumps as by Auchtung and colleagues (Auchtung et al., 2016). These waste lines were connected to a stopcock (Female Luer 4 way, SCLP-400E, Component Supply Co, USA) connected to a 24 deep well sample collection plate sealed with a pierceable sealing mat (AM-24-SQ, AYXMAT, USA) located within the anoxic chamber and to waste bottles located outside of the chamber. Pumps were set to 2.25 for media in and 4.5 for media out.

### Media

The small intestinal media was comprised of the base medium which consists of 0.3 g/L ammonium sulfate, 0.25 g/L sodium chloride, 0.125 g/L L-alanine, 0.095 g/L L-arginine, 0.105 g/L glycine, 0.035 g/L L-histidine, 0.09 g/L L-isoleucine, 0.135 g/L L-leucine, 0.12 g/L L-lysine monochloride, 0.03 g/L L-methionine, 0.085 g/L L-phenylalanine, 0.09 g/L L-proline, 0.045 g/L L-serine, 0.04 g/L DL-threonine, 0.1 g/L valine, 1 ml 0.5% hemin, 1 ml 1% magnesium sulfate, 1 ml 1% calcium chloride, 1 ml 1% sodium sulfate, 2 ml tween 80, and 1 ml trace mineral mix as in Auchtung et al. (Auchtung et al., 2016) with the alteration of 100 mg of nicotinic acid. The supplement consists of 0.3 g/L inulin, 0.287 g/L glutamic acid sodium salt, 16 mg/L asparagine, 9.5 mg/L cysteine, 2.5 mg/L glutamine, 12 mg/L tryptophan, 22.5 mg/L tyrosine, 25 mg/L aspartic acid, 4% potassium phosphate dibasic, 4% potassium phosphate monobasic, 1 ml trace Vitamin Mix as in Auchtung et al. (Auchtung et al., 2016), and 0.5% vitamin K3. This mixture was pH adjusted to 7.2, after which we added 2 g/L sodium bicarbonate. The bile mixture was made of 24.7 mg/L sodium taurocholate hydrate, 45.3 mg/L sodium glycocholate hydrate, 24 mg/L sodium taurochenodeoxycholate, and 43.9 mg/L sodium glycochenodeoxycholate. Sucrose, HFCS (5.5 grams fructose, 4.5 grams glucose), or trehalose were added at 10 g/L. All concentrations are given as final concentrations. Please see Supplemental Table 2 for further details on how to prepare the media.

### Strains

All strains and bioreactors were maintained under anoxic conditions (5% H^2,^ 5% CO_2_, 90% N_2_) at 37ºC. Strains used are shown in Table 1. *Prevotella histicola* DSM 26979 was obtained from the DSMZ culture collection. *Lacticaseibacillus casei* was obtained from a human fecal sample. *Limosilactobacillus reuteri* ATCC PTA 6475, a breast milk isolate, was obtained from BioGaia (Sweden). *Streptococcus mitis* was isolated from breast milk. *Streptococcus salivarius* was isolated from a human fecal sample. *Veillonella atypica* DSM 20739 was obtained from the DSMZ culture collection. All strains were grown on LYBHI (Miquel et al., 2016) plates. All strains were grown in liquid LYBHI except for *P. histicola* which was grown in chopped meat carbohydrate broth (AG20H, Hardy Diagnostics, USA), and *V. atypica* which was grown in BHIS with 0.6% sodium lactate. All strains were sequence confirmed by full-length Sanger sequencing of the 16S rRNA gene. Conditions for these polymerase chain reactions (PCRs) were as follows: a very small bit of microbial colony was added to a 20 µl reaction consisting of 10 μl of Choice Taq (Thomas Scientific, FisherScientific, USA) and 100 nM of each primer (8F and 1492R, **Suppl Table 3**); Cycling conditions were 94ºC for 10 mins, 35 cycles of 94ºC for 45 secs, 50ºC for 1 min, 72ºC for 1 min, followed by a final extension at 72ºC for 10 mins. PCRs were treated with ExoSAP-IT PCR Product Cleanup Reagent (Applied Biosystems, USA) in 5 μl reactions with 1.5 μl of PCR product and 0.25 μl of water following the manufacturer’s reaction condition recommendations. Purified product was mixed with the appropriate primer and sent for Sanger sequencing at GENEWIZ (USA). Diagnostic primers were also utilized to rapidly determine the presence/absence of each strain via gel-electrophoresis. Reaction conditions were similar except for an annealing temperature of 58ºC and a final extension time of 8 mins. See Supplemental Table 3 for all primer sequences.

### Bioreactor run

Twelve reactors were filled with approximately 5 ml of sucrose small intestinal media and rested for 1 day to confirm sterility. Reactors were then inoculated with 500 μl of a stationary phase culture of *P. histicola*. Between 24 and 36 hours later, reactors were inoculated with 500 μl of *L. casei, S. mitis, S. salivarius, L. reuteri* mixed in equal OD_600_ corrected volumes (OD_600_ values were between 0.2 and 0.4) to an approximate OD_600_ of 0.0625 each or 0.25 total. The peristaltic pumps were turned on to 0.5 in and 1 out to allow the reactors to slowly fill overnight. Daily, beginning the following morning, sampling of the bioreactors was achieved by switching the stopcock to flow reactor communities into a 24-well collection plate plated on ice in the anoxic chamber. Sampling was allowed to go for 1.5 to 2 hrs. Following sampling, pumps were increased to 2.25 in and 4.5 out. Two days later, 50 μl of a stationary culture *V. atypica* was inoculated into the reactors. Outflowing media volumes were measured periodically to ensure all reactors were flowing at the same rate, and the occlusion knobs on the peristaltic pump were adjusted as needed. After ∼50 generations (day 6 since the flow rate was set), media in four of the reactors were changed to HFCS small intestinal media, four to trehalose small intestinal media, while the other four were maintained on sucrose small intestinal media. After ∼100 more generations (day 18), the pumps were stopped and the cultures growing within the reactors were harvested. Periodically, sampled cultures were plated in serial dilutions on LYBHI plates for colony counting. Cultures from generations 50, 100, and 150 were also saved mixed with 10% DMSO to preserve live microbial communities. Generations are calculated as time past in hours * number of reactor turnovers per hour * 1/ln(2) as previously described (Van den Bergh et al., 2018). See Supplemental Figure 6A for an overview of the bioreactor run. Sequencing of the inoculates suggested that the mixture of *L. casei, S. mitis, S. salivarius, L. reuteri* inoculate was biased towards *S. mitis* and *S. salivarius* (**Suppl Fig 6B**).

### Community composition analysis

DNA was isolated from microbial community cell pellets using the DNeasy Blood & Tissue kit (Qiagen, USA). DNA was quantified using the Quant-iT dsDNA Assay kit high sensitivity kit (ThermoFisher, USA). Microbial community composition was determined by Illumina sequencing of the V4 region of the 16S rRNA gene followed by analysis using QIIME 2 2019.7 (Bolyen et al., 2019). 16S rRNA V4 amplicon libraries were prepared using 25 µg of DNA in two 50 µl PCRs using Phusion High-Fidelity DNA Polymerase (ThermoFisher, USA) with the following reaction conditions: 1x Phusion HF Buffer, 0.5 µM of each forward and reverse primer (see Supplemental Table 3), 200 µM dNTPs, and 0.01 U/μl Phusion High-Fidelity DNA Polymerase. These PCRs were run in a thermocycler with the following conditions: 98ºC for 30 secs, 30 cycles of 98ºC for 10 secs, 53ºC for 15 secs, 72ºC for 20 secs, followed by 72ºC for 5 mins. Libraries were purified using AMPure XP beads (Beckman Coulter, USA), quantified using Quant-iT dsDNA Assay high sensitivity kit, and pooled to 100 µg. The library was submitted along with sequencing primers (see Supplemental Table 3) to the Microbial Genome Sequencing Center (MiGS, Pittsburgh, PA, USA).

For QIIME analysis, demultiplexed data provided by MiGS were denoised using DADA2 (Callahan et al., 2016) with the options trim-left-r 5, trunc-len-f 240, trunc-len-r 206. Representative sequences were aligned in a pairwise blastn (Altschul et al., 1990) to the 16S rRNA genes of our six microbes. Counts for representative sequences matching the same microbe at >98% identity and 100% query coverage were merged. Several low abundance OTUs assigned to Blautia and Bifidobacteria that were only present in the day 3 samples and thus likely represent a contaminant introduced in the taken sample were removed. Finally, we removed singlet OTUs appearing in a single sample with counts <10. OTU counts were rarefied to 2413. Percent abundances of each microbe were corrected for 16S rRNA gene copy number differences. Copy numbers used were 5 for *L. casei*, 6 for *L. reuteri*, 4 for *S. mitis*, 6 for *S. salivarius*, 3 for *V. atypica*, and 4 for *P. histicola*.

Principal coordinate analysis was performed using the distance matric from the Bray Curtis dissimilarity metric (Bray and Curtis, 1957) in the phyloseq package (McMurdie and Holmes, 2013). Significance of separation of the reactors by sugar was computed using a PERMANOVA in adonis in the vegan package (Oksanen et al., 2019). Differential abundance of strains before and after the sugar media change was determined by first performing a Kruskal-Wallis test on each microbe’s abundance per sugar media on time points before and after the media change. Resulting p-values were adjusted using a Benjamini-Hochberg correction (Benjamini and Hochberg, 1995), with adjusted p-values <0.1 considered significant. For microbes reporting significance, which media comparison was significant was computed using Dunnett’s Test with sucrose set as the control implemented using the DunnettTest function in DescTools (Andri et mult. al. S, 2021) with p <0.05 considered significant.

Species evenness was assessed using the renyiresult function in BiodiversityR (Kindt and Coe, 2005). Significance of these diversity metrics among the reactors was determined using Dunnett’s Test with sucrose set as the control implemented using the DunnettTest function in DescTools (Andri et mult. al. S, 2021).

### Growth curves

Communities from generations 50, 100, and 150 for each reactor were regrown from a scraping of their DMSO stock into 200 µl of the media the community was grown in post generation 50 for two days. Then the culture was spun down, washed in an equal volume of 0.85% NaCl, and diluted 1/10 into a total of 200 µl of each of the different sugar media in a 96-well plate. The culture growth was measured by taking OD_600_ measurements every 10 minutes for 16.5 hours on a continually shaking plate reader. Each community was grown in triplicate and the entire assay was repeated. ODs from averaged blank wells were subtracted from the data for their respective media. Blanked data were averaged across all replicates. Growth assays were conducted at 5% H_2,_ 5% CO_2_, and 90% N_2_. Growth curves for individual microbial strains in each sugar media were similarly obtained.

### Organoid maintenance

Human jejunal *NGN3* enteroendocrine cell enriched organoids were maintained as previously described (Chang-Graham et al., 2019). Briefly, organoids were embedded in Matrigel (Corning, USA) and cultured in complete media with growth factors (CMGF+) supplemented with 50% (v/v) Wnt3a conditioned medium, 10 µmol Y-27632 Rock inhibitor (InvivoGen, USA), and 200 µg/ml Geneticin (Gibco, USA) in a humidified 5% CO_2_ incubator at 37ºC. Organoids were routinely passaged every 7 days and less than 50 times in total.

### Hormonal secretion assay

Organoid monolayers were produced by coating 96 well plates in 25 µl/ml Matrigel and seeding them with two 3D wells of *NGN3* organoids per one 96 well. After two days recovery in CMGF+, monolayers were differentiated using differentiation medium (Chang-Graham et al., 2019) supplemented with Rock inhibitor and 1 µg/ml doxycycline. After 5 days differentiation, media were removed and replaced with pH neutralized and 0.22 µm filtered bioreactor supernatant that had been lyophilized and resuspended without dilution into Krebs buffer (Sigma-Aldrich, USA). Monolayers were incubated in a humidified 5% CO_2_ incubator at 37ºC for 3 hours. Following, supernatants were removed and frozen at −20ºC. Serotonin was measured using an ELISA (Eagle Biosciences, USA) and other hormones using the Luminex MILLIPLEX Human Metabolic Hormone kit (EMD Millipore, USA). Differences in hormone concentrations or fold changes were determined first with a Kruskal Wallis test and where significant, a Dunn’s test with a Benjamini-Hochberg multiple comparison adjustment was used to obtain the significance of individual comparisons, implemented using the DunnTest function in DescTools (Andri et mult. al. S, 2021).

### Linear modeling

For hormones with differential secretion across the sugar small intestinal media, we determined the contribution of each microbe and media type to the secreted amount using linear modeling. We excluded one of the organoid monolayer replicates from sucrose reactor 1 that gave low secretion data for many of the measured hormones. Hormone data from organoids treated with media controls were included with the fractional abundance of each microbe set to 0. For each hormone, we began with a full model of the form: *Concentration ∼ (Lcas + Lreu + Phis + Smit + Ssal + Vaty) * media* where *Lcas, Lreu, Phis, Smit, Ssal*, and *Vaty* are the fractional abundance of *Lactobacillus casei, Lactobacillus reuteri, Prevotella jejuni, Streptococcus mitis, Streptococcus salivarius*, and *Veillonella atypica* respectively derived from copy number corrected 16S rRNA sequencing data from the reactors at the end of the bioreactor experiment. Microbes were removed sequentially if removal did not significantly affect the fit of the model as assessed by a likelihood ratio test. For plotting, interactions that were reported as significant (p<0.05) in the resulting models were plotted as regression lines from the reported intercepts and coefficients. Bounding lines were also plotted from the standard errors to create a shaded area showing the error.

### Statistical analyses and plotting tools

All statistical analyses were conducted in R (R Core Team, 2021). Graphs were made using ggplot2 (Wickham, 2016). P<0.05 and p-adjust<0.1 were considered significant.

## Data availability

Illumina sequencing reads of the bioreactor microbial communities are available in the European Nucleotide Archive under accession PRJEB54023. Raw data and code used to generate figures are available at https://github.com/sdirienzi/CAMIO.

## Legends

**Supplemental Figure 1: Small intestinal bioreactor community composition and density. A)** Principal coordinate analysis on a Bray-Curtis distance dissimilarity matrix of each bioreactor’s microbial community for each sampled generation/day. All data are shown in each plot with colored points representing the named generation/day. Points are colored by bioreactor sets (connected to the same media bottle). When differences among the sets were significant by PERMNOVA, the F-stat, R^2^, and Benjamini-Hochberg adjusted p-value (over all time points) are shown as well as ellipses are drawn using a t-distribution. **B)** Cell density per ml in the reactors as assessed by plating on LYBHI plates. Shape indicates individual reactors for each media type: reactor 1 = circle; reactor 2 = square; reactor 3 = diamond; reactor 4 = triangle. Generations are averaged across reactors.

**Supplemental Figure 2: Microbial community composition in response to different sugars**. Community composition of reactor flow-through throughout the run. The fraction of each microbe determined by 16S rRNA sequencing is shown: *L. casei* (blue), *L. reuteri* (orange), *P. histicola* (green), *S. mitis* (red), *S. salivarius* (purple), *V. atypica* (brown). The gray boxes indicate when the reactors were on sucrose media before being changed to HFCS or trehalose media.

**Supplemental Figure 3: Microbial community growth adaptation in different sugars**. Growth dynamics of each reactor community at generation 50, 100, or 150 when regrown in batch culture in each sugar media. Lines are colored by media type: sucrose as purple; HFCS as orange; trehalose as pink. Line type indicates from which generation the community was derived: 50 as dotted; 100 as dashed; and 150 as solid.

**Supplemental Figure 4: Hormones and cytokines secreted from human intestinal organoids treated with supernatants from the microbial communities**. Concentration of **A)** C-peptide, **B)** ghrelin, **C)** GIP, **D)** GLP-1, **E)** MCP-1, **F)** PP, **G)** PYY, **H)** serotonin, and **I)** TNFα from enteroendocrine-enriched organoid monolayers treated with bioreactor supernatants. Reactors are named as R1 through R4 for each sugar media. Media controls are labeled M1 and M2 and differ only in the batch of organoids they were applied to. A single batch of organoids was used for reactors R1 through R3 and media control M1, and a single batch of organoids was used for reactors R4 and media control M2. The gray line indicates the assay’s lower limit of detection. Data from triplicate organoid monolayers are shown for each supernatant treatment.

**Supplemental Figure 5: Growth dynamics of individual small intestinal strains on small intestinal media**. Growth of **A)** *L. casei*, **B)** *L. reuteri*, **C)** *L. salivarius*, **D)** *L. mitis*, **E)** *P. histicola*, **F)** *V. atypica* growing on sucrose, HFCS, or trehalose small intestinal media. Vertical lines indicate the standard deviation determined from triplicates.

**Supplemental Figure 6: Small intestine bioreactor run**. Bioreactor design: **A)** Twelve bioreactors were filled with sucrose small intestinal media and inoculated with *P. histicola* two days prior to starting flow. The following day, reactors were inoculated with *S. salivarius, S. mitis, L. casei*, and *L. reuteri*. The next day, the flow rate was set to a 3.3 hr turnover and daily sampling began. Two days later, reactors were inoculated with *V. atypica*. After 50 generations (6 days), four reactors were switched to HFCS small intestinal media and four to trehalose small intestinal media. The reactor run continued for 100 more generations. **B)** Composition of the inoculates. The fraction of each microbe determined by 16S rRNA sequencing is shown for each inoculate: *L. casei* (blue), *L. reuteri* (orange), *P. histicola* (green), *S. mitis* (red), *S. salivarius* (purple), *V. atypica* (brown).

**Supplemental Table 1: Linear modeling of hormone concentrations as a function of microbial abundances and media type**

**Supplemental Table 2: Small Intestinal Media Supplemental Table 3: Primers**

