## Supplementary figures and images for "Separation of the effects of small intestinal microbiome-diet interactions on human gut hormone secretion"

### Supplemental Figure 1

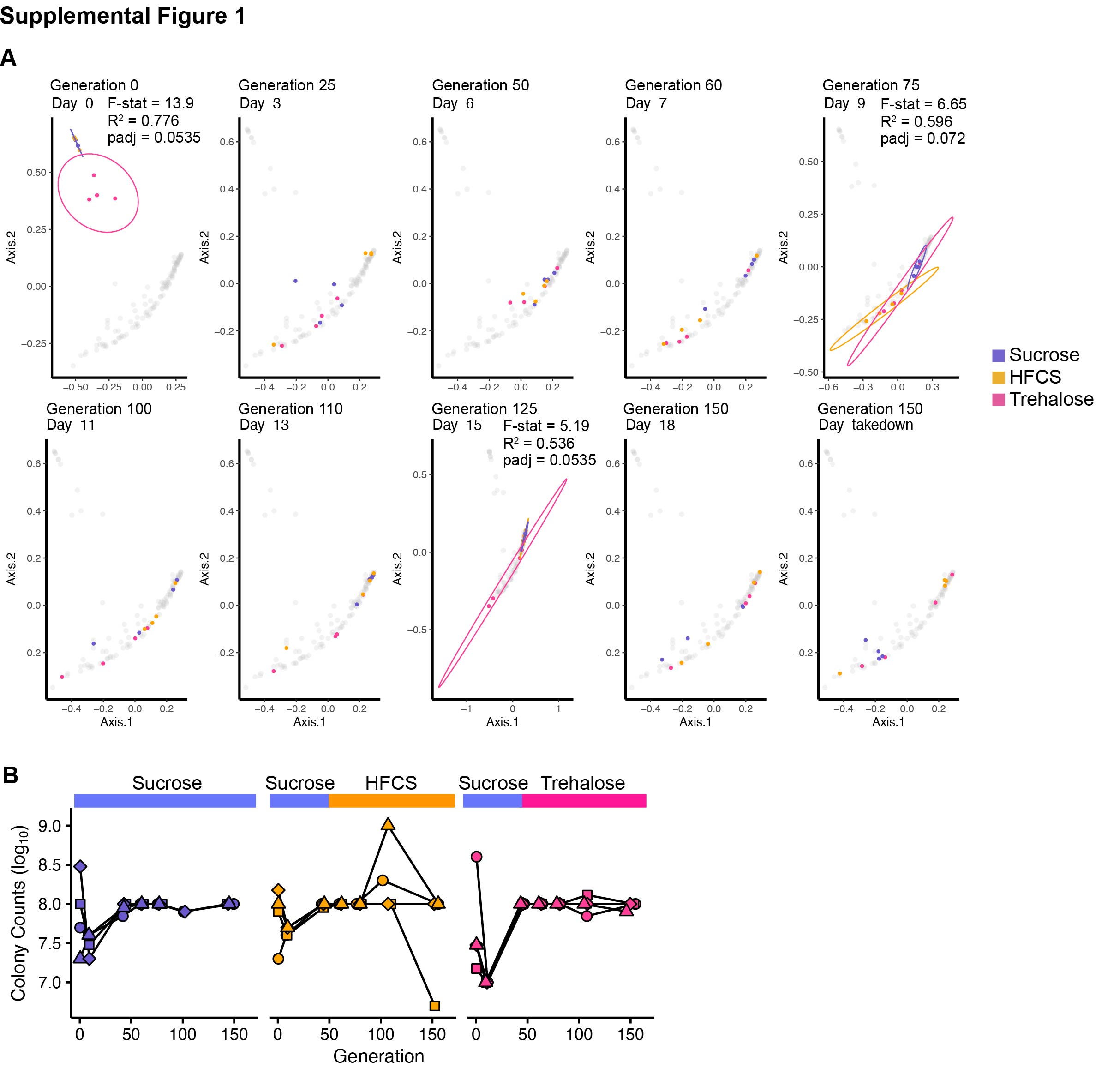

### Supplemental Figure 2

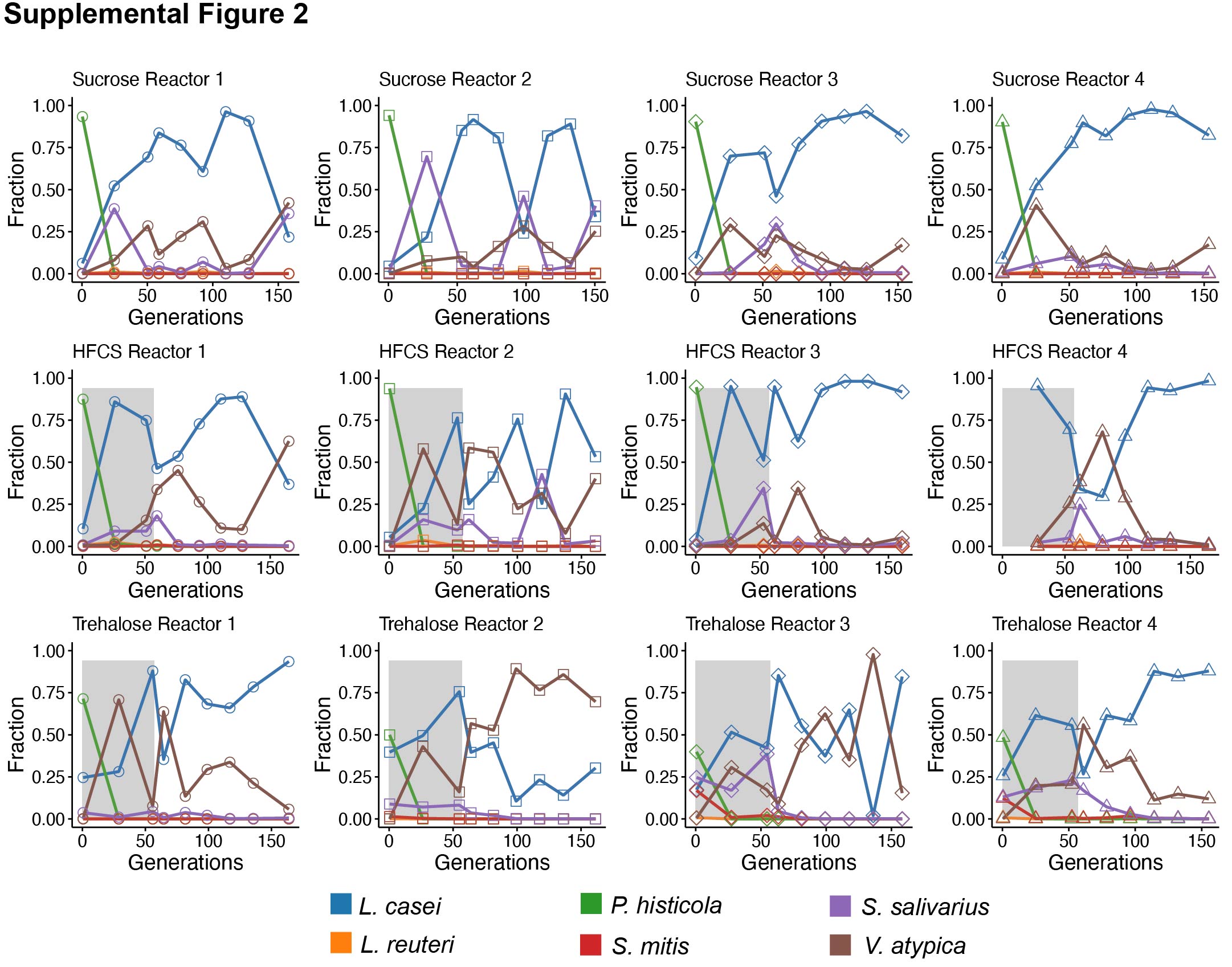

### Supplemental Figure 3

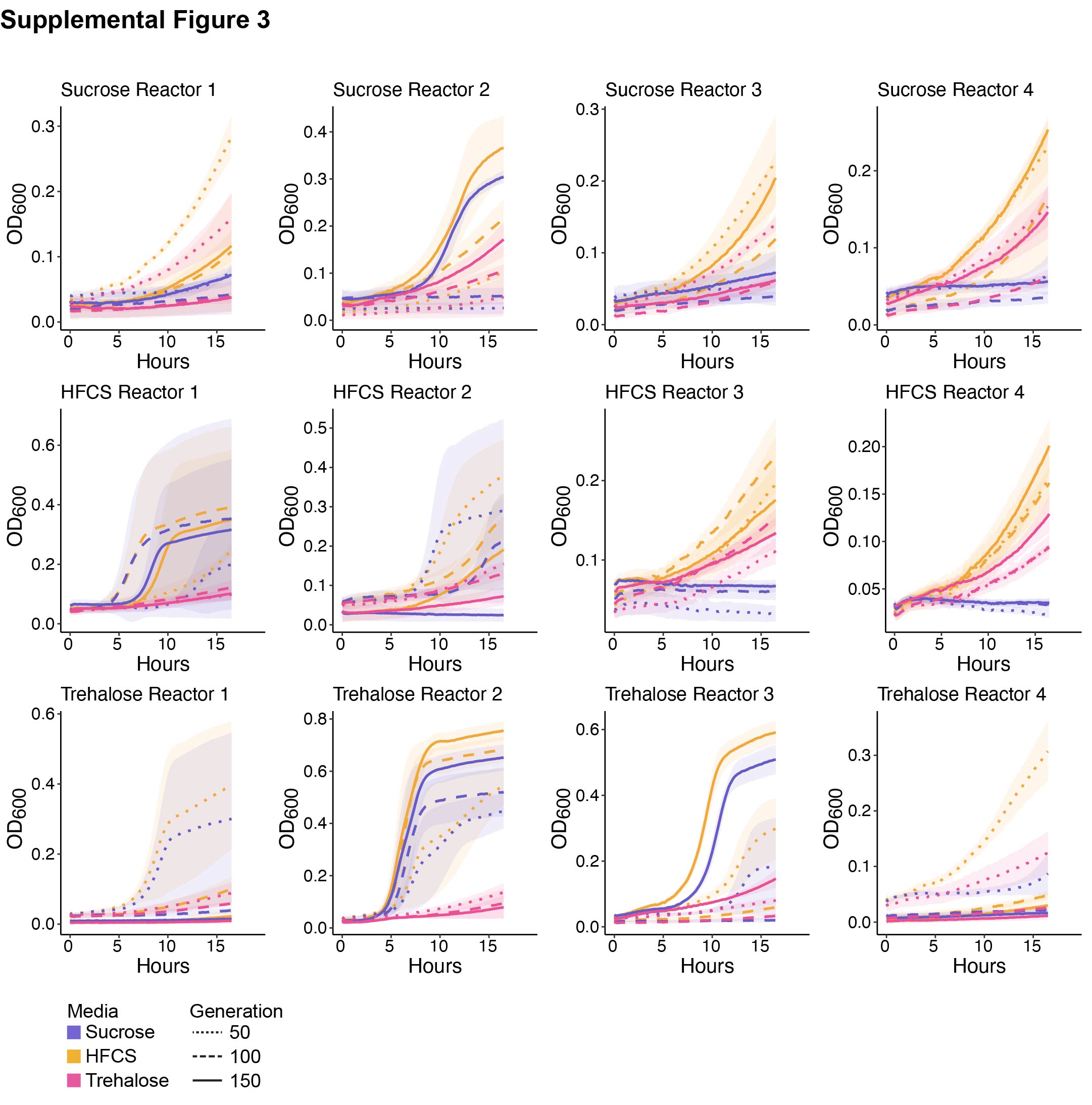

### Supplemental Figure 4

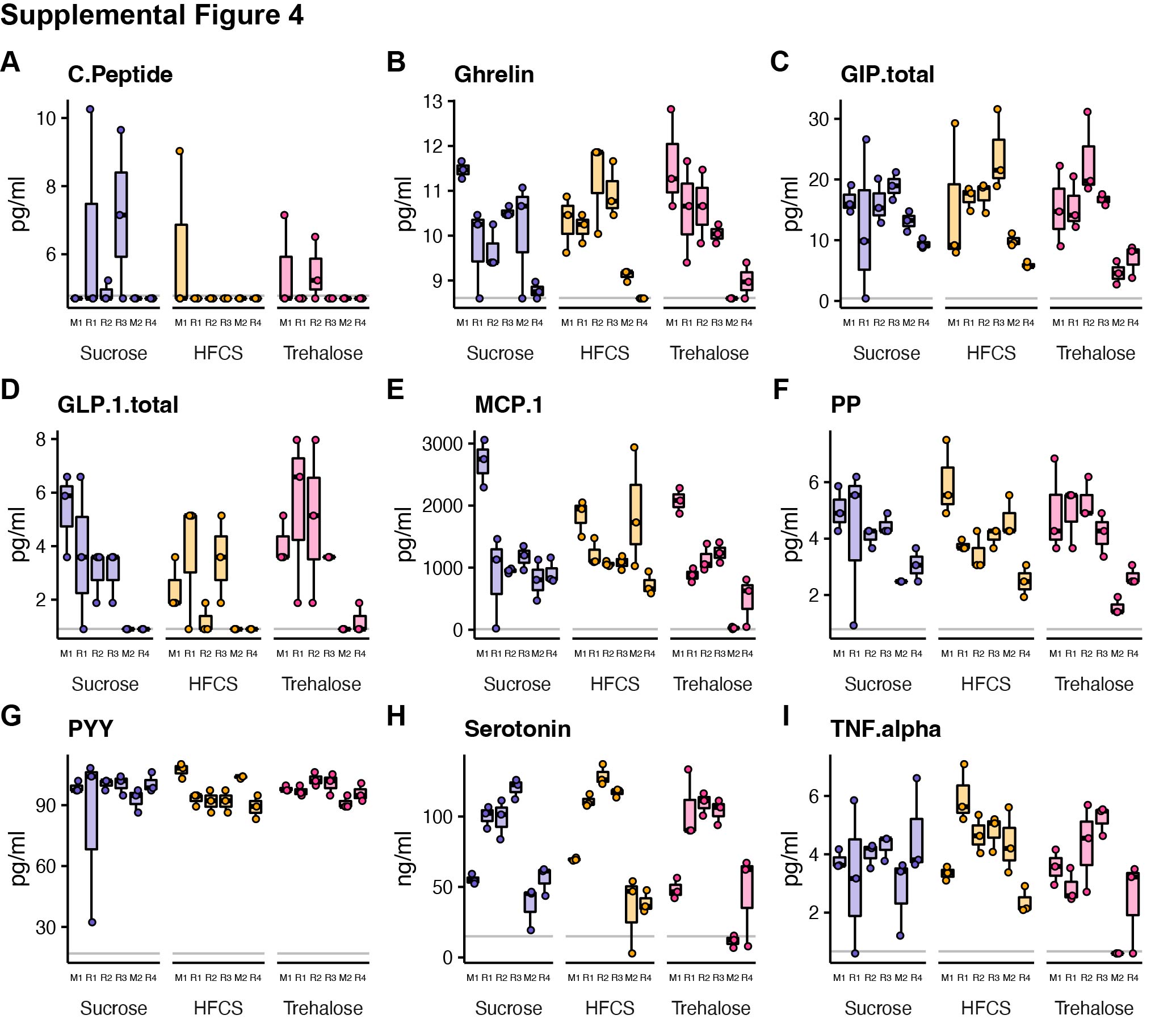

### Supplemental Figure 5

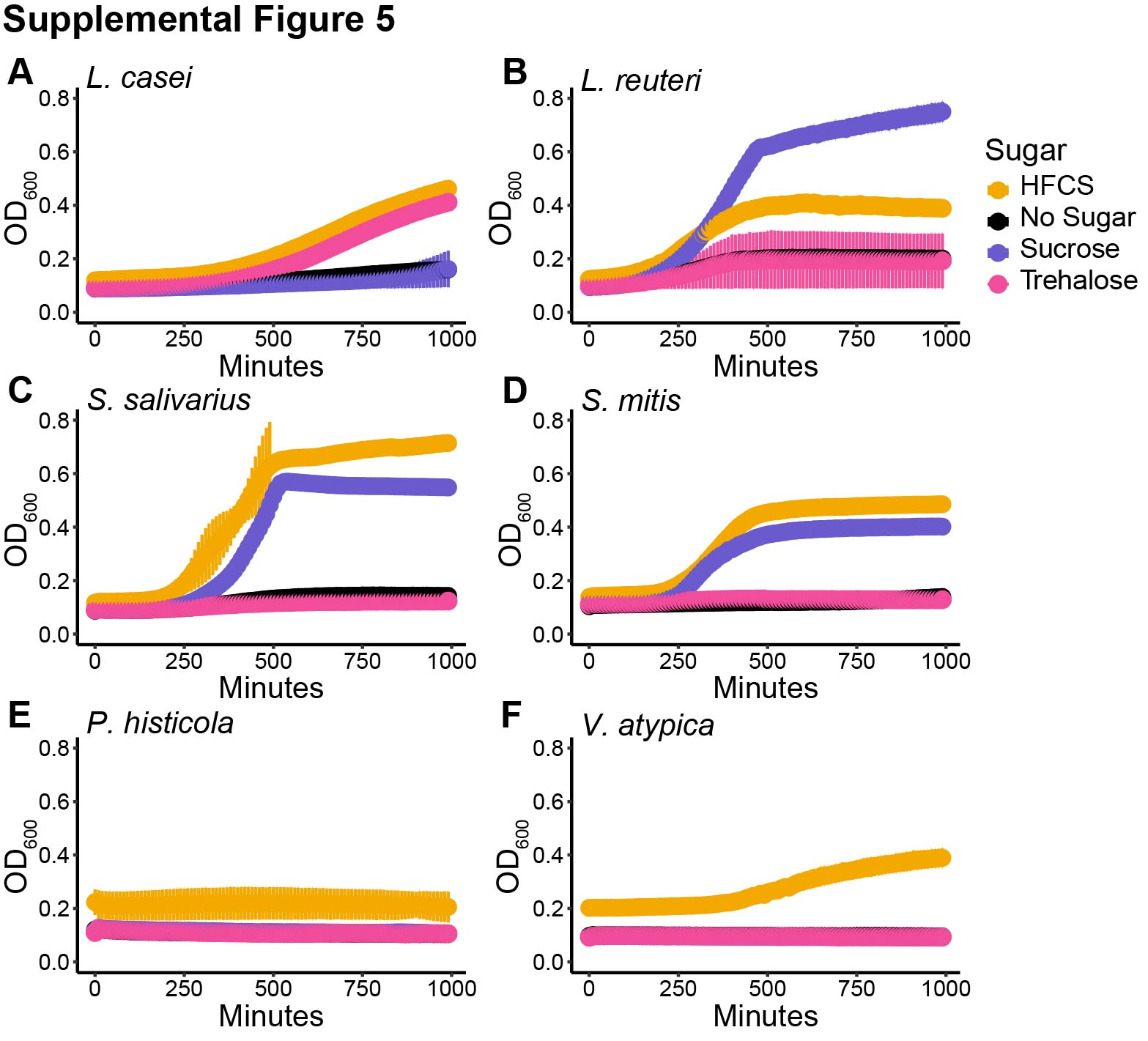

### Supplemental Figure 6

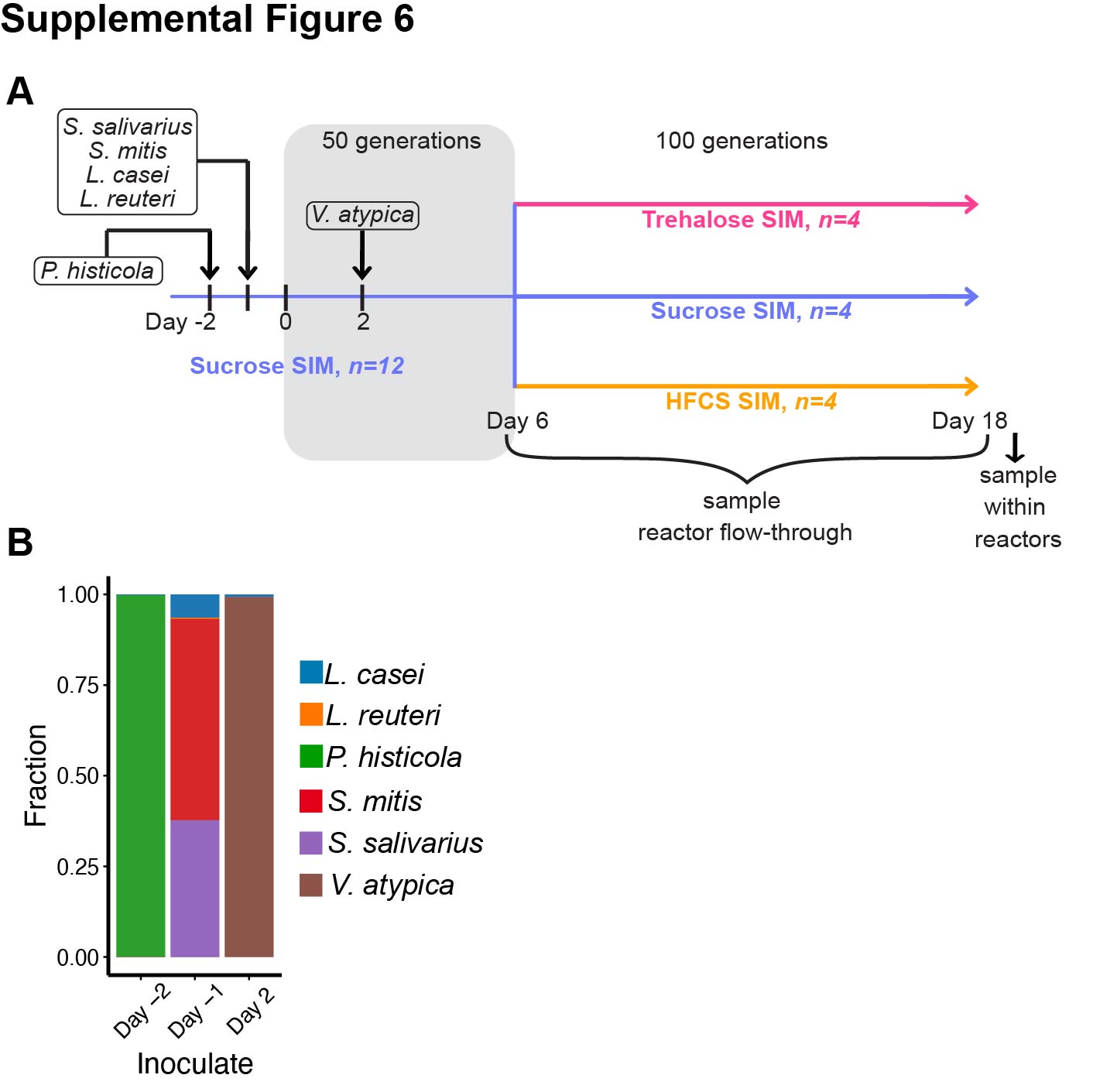
