## Supplemental Tables for "Separation of the effects of small intestinal microbiome-diet interactions on human gut hormone secretion"

**Supplemental Table 1: Linear modeling of hormone concentrations as a function of microbial abundances and media type**

**Ghrelin: Concentration~(Lcas + Ssal + Vaty)*Media**

| Term | Estimate | Std Error | t value | P-value |
|---|---|---|---|---|
| Intercept (Sucrose) | 10.7791 | 0.3838 | 28.087 | <2.00E-16 |
| Lcas | 17.0682 | 8.6725 | 1.968 | 0.05585 |
| Ssal | -2.4824 | 1.4848 | -1.672 | 0.10215 |
| Vaty | -14.7739 | 6.4944 | -2.275 | 0.02821 |
| HFCS | -1.0561 | 0.5428 | -1.946 | 0.05857 |
| Trehalose | -0.6878 | 0.5428 | -1.267 | 0.21225 |
| Lcas:HFCS | -14.3821 | 8.7087 | -1.651 | 0.10628 |
| Lcas:Trehalose | -18.0426 | 8.6959 | -2.075 | 0.04432 |
| Ssal:HFCS | -24.8297 | 7.8666 | -3.156 | 0.00299 |
| Ssal:Trehalose | 24.8702 | 13.2203 | 1.881 | 0.06706 |
| Vaty:HFCS | 16.7228 | 6.5543 | 2.551 | 0.01455 |
| Vaty:Trehalose | 14.4937 | 6.5822 | 2.202 | 0.03335 |

Residual standard error: 0.9403; Multiple R-squared: 0.4132; F-statistic: 2.624; p-value: 0.01236

**PP: Concentration~Ssal*Media**

| Term | Estimate | Std Error | t value | P-value |
|---|---|---|---|---|
| Intercept (Sucrose) | 4.0258 | 0.341 | 11.805 | 1.16E-15 |
| Ssal | 0.2871 | 1.5695 | 0.183 | 0.855667 |
| HFCS | 1.0732 | 0.5219 | 2.056 | 0.045347 |
| Trehalose | -0.8751 | 0.4978 | -1.758 | 0.085254 |
| Ssal:HFCS | -23.8426 | 6.7618 | -3.526 | 0.000953 |
| Ssal:Trehalose | 38.4643 | 11.9444 | 3.22 | 0.002325 |

Residual standard error: 1.158; Multiple R-squared: 0.3356; F-statistic: 4.747; p-value: 0.001363

**PYY: Concentration~(Lcas+Vaty)*Media**

| Term | Estimate | Std Error | t value | P-value |
|---|---|---|---|---|
| Intercept (Sucrose) | 95.819 | 1.695 | 56.533 | <2.00E-16 |
| Lcas | 18.438 | 33.998 | 0.542 | 0.59034 |
| Vaty | -1.513 | 25.815 | -0.059 | 0.953515 |
| HFCS | 9.55 | 2.429 | 3.931 | 0.000296 |
| Trehalose | -1.212 | 2.43 | -0.499 | 0.620277 |
| Lcas:HFCS | -34.301 | 34.091 | -1.006 | 0.319832 |
| Lcas:Trehalose | -17.513 | 34.106 | -0.513 | 0.610171 |
| Vaty:HFCS | -11.178 | 26.067 | -0.429 | 0.670146 |
| Vaty:Trehalose | 12.582 | 26.078 | 0.482 | 0.631844 |

Residual standard error: 4.266; Multiple R-squared: 0.5963; F-statistic: 8.123; p-value: 1.15e-06

**Supplemental Table 2: Small Intestinal Media**

| Base medium | | | |
|---|---|---|---|
| Ingredient | mass/volume | source | product number |
| $diH_2O$ | 100 ml | NA | NA |
| Ammonium sulfate | 0.3 g | MP Biomedicals, LLC (Solon, OH, USA) | 150373 |
| Sodium chloride | 0.25 g | Fisher Chemical (Fair Lawn, NJ, USA) | S271-3 |
| L-alanine | 0.125 g | Sigma Aldrich (St. Louis, MO, USA) | A7469 |
| L-arginine | 0.095 g | Acros Organics (Morris Plains, NJ, USA) | 104881000 |
| Glycine | 0.105 g | Sigma Aldrich (St. Louis, MO, USA) | G7126 |
| L-histidine | 0.035 g | Alfa Aesar (Shore Rd, Heysham, England) | A10413 |
| L-isoleucine | 0.09 g | Calbiochem (Billerica, MA, USA) | 4160 |
| L-leucine | 0.135 g | Sigma Aldrich (St. Louis, MO, USA) | L8000 |
| L-lysine | 0.12 g | Acros Organics (Morris Plains, NJ, USA) | 125221000 |
| L-methionine | 0.03 g | Fisher BioReagents (Fair Lawn, NJ, USA) | BP388 |
| L-phenylalanine | 0.085 g | Alfa Aesar (Shore Rd, Heysham, England) | A13238 |
| L-proline | 0.09 g | Sigma Aldrich (St. Louis, MO, USA) | P0380 |
| L-serine | 0.045 g | Sigma Aldrich (St. Louis, MO, USA) | S4500 |
| DL-threonine | 0.04 g | Chem-Impex Int'l Inc (Wood Dale, IL, USA) | 00368 |
| Valine | 0.1 g | Alfa Aesar (Shore Rd, Heysham, England) | A12720 |
| 0.5% Hemin in 1.4M NaOH | 1 ml | Frontier Science (Logan, UT, USA) | H651-9 |
| 1% Magnesium sulfate | 1 ml | Fisher BioReagents (Fair Lawn, NJ, USA) | BP213-1 |
| 1% Calcium chloride | 1 ml | FisherBiotech (Fair Lawn, NJ, USA) | BP510 |
| 1% Sodium sulfate | 1 ml | Sigma Aldrich (St. Louis, MO, USA) | 239313 |
| Tween 80 | 2 ml | VWR Life Sciences (Solon, OH, USA) | 0442 |
| Trace mineral mix | 1 ml | see right | see right |
| filter sterile store at 4C | | | |

| Supplement | | | |
|---|---|---|---|
| Ingredient | mass/volume | source | product number |
| $diH_2O$ | 40 ml | NA | NA |
| Inulin | 0.3 g | Alfa Aesar (Shore Rd, Heysham, England) | A18425 |
| Dissolve inulin in H20. Heat gently until in solution | | | |
| Remove from heat and continue stirring til dissolved, then add | | | |
| L-glutamic acid monosodium salt | 0.287 g | Sigma Aldrich (St. Louis, MO, USA) | G1626 |
| L-asparagine | 16 mg | Sigma Aldrich (St. Louis, MO, USA) | A0884 |
| L-cysteine | 9.5 mg | Sigma Aldrich (St. Louis, MO, USA) | C7532 |
| L-glutamine | 2.5 mg | FisherBiotech (Fair Lawn, NJ, USA) | BP379 |
| L-tryptophan | 12 mg | Sigma Aldrich (St. Louis, MO, USA) | T0254 |
| 4% Potassium phosphate dibasic anhydrous | 1 ml | Fisher Chemical (Fair Lawn, NJ, USA) | P288 |
| 4% Potassium phosphate monobasic | 1 ml | Fisher BioReagents (Fair Lawn, NJ, USA) | BP362 |
| Trace vitamin mix | 1 ml | see right | see right |
| 0.5% Vitamin $K_3$ in ethanol | 0.2 ml | Enzo Life Sciences (Farmingdale, NY, USA) | 460-007 |
| L-tyrosine | 22.5 mg | Sigma Aldrich (St. Louis, MO, USA) | T3754 |
| L-aspartic acid | 25 mg | Acros Organics (Morris Plains, NJ, USA) | 105041000 |
| Adjust pH to 7.2 with NaOH, then add | | | |
| Sodium bicarbonate | 2 g | Fisher Chemical (Fair Lawn, NJ, USA) | S233 |
| filter sterile and store at 4C | | | |

| Bile | | | |
|---|---|---|---|
| Mix together with no heat | | | |
| Ingredient | mass/volume | source | product number |
| $diH_2O$ | 1 | NA | NA |
| Sodium taurocholate hydrate | 0.0247 g | Sigma Aldrich (St. Louis, MO, USA) | T4009 |
| Sodium glycocholate hydrate | 0.0453 g | Sigma Aldrich (St. Louis, MO, USA) | G7132 |
| Sodium taurochenodeoxycholate | 0.0240 g | Sigma Aldrich (St. Louis, MO, USA) | T6260 |
| Sodium glycochenodeoxycholate | 0.0439 g | Sigma Aldrich (St. Louis, MO, USA) | G0759 |
| filter sterilize through a syringe filter, store at room temp | | | |

| 1L Trace Vitamin Mix (1000X) | |
|---|---|
| Folic acid | 15 mg |
| Pyridoxine hydrochloride | 60 mg |
| Riboflavin 5 phosphate sodium salt hydrate | 213 mg |
| Biotin | 2 mg |
| Thiamine hydrochloride | 58 mg |
| Nicotinic acid | 100 mg |
| Calcium pantothenate | 120 mg |
| Vitamin B12 | 1 mg |
| p-Aminobenzoic acid | 12 mg |
| Lipoic acid | 5 mg |
| Potassium Phosphate | 900 mg |
| ph to 7 with NaOH and filter | |

use sterile water, no autoclave or filter

| 1L Trace Mineral Mix (1000X) | |
|---|---|
| Hydrochloric Acid | 1.6 ml |
| $FeSO_4.7H_2O$ | 2.1 g |
| $H_3BO_3$ | 30 mg |
| $MnCl_2.4H_2O$ | 100 mg |
| $CoCl_2.6H_2O$ | 190 mg |
| $NiCl_2.6H_2O$ | 24 mg |
| $CuCl_2.2H_2O$ | 2 mg |
| $ZnSO_4.7H_2O$ | 144 mg |
| $Na_2MoO_4.2H_2O$ | 36 mg |
| $NaVO_3$ | 25 mg |
| $Na_2WO_4.2H_2O$ | 25 mg |
| $Na_2SeO_3.5H_2O$ | 6 mg |

| Sugar | | | |
|---|---|---|---|
| Add one of the following to 10 mls of diH2O with no heat | | | |
| Sucrose | 10 g | Fisher BioReagents (Fair Lawn, NJ, USA) | BP220-1 |
| Fructose | 5.5 g | Alfa Aesar (Ward Hill, MA, USA) | A17718 |
| Glucose | 4.5 g | VWR Chemical BDH (Radnor, PA, USA) | BDH9230 |
| Trehalose, dihydrate | 11.05 | TCI America (Portland, OR, USA) | T0331 |

To mix the component togethers per bottle
Measure out the base media

Add diH2O
Filter sterilize into a sterile bottle with a sterile stir bar
Filter sterilize other components into the same bottle
Mix with stirring

| | for 1 liter |
|---|---|
| base medium | 106 ml |
| filter sterilized water | 795 ml |
| supplement | 48 ml |
| bile | 1 ml |
| sugar | 50 ml |
| total | 1000 ml |

**Supplemental Table 3: Primers**

| Primer Names | Forward Sequence | Reverse Sequence | Annealing Temperature | Fragment Size | Purpose | Citation |
|---|---|---|---|---|---|---|
| 8F, 1492R | AGAGTTTGATCCTGGCTCAG | GGTTACCTTGTTACGACTT | 50 | 1484 | Amplify whole 16SrRNA gene | Edwards et al. 1989. Nucleic Acids Research 17: 7843-7853; Stackebrant and Liesack. 1993. Nucleic acids and classification, p. 152-189. |
| PhisF, PhisR | GGAAACGGCATTAAGTGCTTGC | ATCCCCATCCATAACCGATAAATC | 58 | 175 | Detect P. histicola | This work |
| LcasF, LcasR | GAGATTCAACATGGAACGAGTGG | CCATCCAAAAGCGATAGCTTACG | 58 | 159 | Detect L. casei | This work |
| LreuF, LreuF | GCTGAAAGACGACGCTTTCTGC | CATTGAACCAGCTGATATTCTTGCTTCC | 58 | 339 | Detect L. reuteri | van Pijeren and Britton. 2012. Nucleic Acids Research 40:e76 |
| SmitF, SmitR | GAAGAATTGCTTGAATTGGTTGAA | GGACGGTAGTTGTTGAAGAATGG | 58 | 559 | Detect S. mitis | Collado et al. 2009. Letters in Applied Microbiology. 48:523-528 |
| SsalF, SsalR | CGTAACGTGGGAAAACTGTCC | GATAACGTTGACCTTACGCTAGC | 58 | 193 | Detect S. salivarius | This work |
| VatyF, VatyR | AYCAACCTGCCCTTCAGA | CGTCCCGATTAACAGAGCTT | 58 | 350 | Detect V. atypica | Rinttila et al. 2004. Journal of Applied Microbiology 97:1166-1177 |
| 16S rRNA 515F, Barcode 1 | AATGATACGGCGACCACCGAGATCTACACATCGATGGTATGGTAATTGGGTGCCAGCMGCCGCGGTAA | | 53 | 350 | Amplify & barcode V4 region of 16SrRNA gene | Auchtung et al. 2015. Microbiome 3:42 |
| 16S rRNA 515F, Barcode 3 | AATGATACGGCGACCACCGAGATCTACACGGTATCTCTATGGTAATTGGGTGCCAGCMGCCGCGGTAA | | 53 | 350 | Amplify & barcode V4 region of 16SrRNA gene | Auchtung et al. 2015. Microbiome 3:42 |
| 16S rRNA 806R | 806rcbc0 to 806rcbc95 | | 53 | 350 | Amplify & barcode V4 | Caporaso et al. 2012. ISME J 6:1621-1624 |
| Read 1 primer | TATGGTAATTGGGTGCCAGCMGCCGCGGTAA | | | | Sequence read 1 on an Illumina MiSeq | Kozich et al. 2013. Appl Environ Microbiol 79:5112-5120 |
| Read 2 primer | AGTCAGTCAGCCGGACTACHVGGGTWTCTAAT | | | | Sequence read 2 on an Illumina MiSeq | Kozich et al. 2013. Appl Environ Microbiol 79:5112-5120 |
| Index primer | ATTAGAWACCCBDGTAGTCCGGCTGACTGACT | | | | Sequence index read on an Illumina MiSeq | Auchtung et al. 2015. Microbiome 3:42 |
